# Prediction of myeloid malignant cells in Fanconi anemia using machine learning

**DOI:** 10.1101/2023.12.31.573791

**Authors:** Luis A. Flores-Mejía, Pablo Siliceo-Portugal, Ulises Juárez Figueroa, Iram García Quezada, Bibiana T. Barcenas Martínez, David Sosa, Anet Rivera, Hugo Tovar, Sara Frías, Alfredo Rodríguez

## Abstract

Fanconi anemia (FA) is an inherited bone marrow failure syndrome with cancer predisposition. Most FA patients develop aplastic anemia during childhood and have an extremely high cumulative risk to develop cancer during their lifespan. Myeloid malignancy is one of the main tumor risks for patients with FA, including high-risk myelodysplastic syndrome (MDS) and acute myeloid leukemia (AML). Although bone marrow transplantation is the treatment of choice for FA patients that develop aplastic anemia, patients with a more stable bone marrow remain at a high risk of presenting MDS/AML and should be monitored for appearance of myeloid malignant clones. Markers for an as-early-as-possible identification of emerging myeloid malignant cells are needed for the monitoring of patients with FA, since quick medical action after detection of neoplastic transformation is needed.

In this work we have leveraged publicly available single cell RNA seq (scRNAseq) datasets of patients with MDS and AML for training deep neural networks (DNN). We have generated two machine learning models aimed to identify myeloid malignant transcriptional profiles in scRNAseq datasets from the bone marrow of patients with FA, one for detection of MDS and a second one for AML. Both predictors displayed high sensitivity, specificity, and accuracy for detection of single cell resolution myeloid malignant transcriptional profiles.

Multiple tools for analysis of single cell transcriptional data were implemented to characterize the predicted MDS and AML cells. Our analysis suggests that the predicted MDS and AML cells from FA patients are enriched in the lympho-myeloid-primed progenitor (LMPP) and the granulocyte-monocyte progenitor (GMP) populations. The predicted MDS and AML cells have gene expression and master transcriptional factor profiles that suggest malignant transformation and that differ from the rest of FA cells. Also cues of immune evasion were detected using single cell pathway analysis (SCPA) and cell-cell communication profiles. Next work will be aimed to find potential cell surface markers on the predicted MDS and AML cells as well as to assess our predictions in primary samples from FA patients.

## BACKGROUND

The inherited bone marrow failure syndromes (IBMFS) are rare entities characterized by physical abnormalities, and an exacerbated risk to present bone marrow failure (BMF)^1,2^. The most recognizable IBMFS is Fanconi anemia (FA). Patients with FA have defects in the FA/BRCA pathway that is responsible for the repair of interstrand crosslinks (ICLs).

Patients with FA have an extremely high predisposition to develop squamous cell carcinomas in the oral cavity and anogenital region^3^, and at the same time have an extreme predisposition to develop myeloid malignancies, specifically myelodysplastic syndrome (MDS) and acute myeloid leukemia (AML).

MDS is a clonal hematopoietic neoplasm characterized by bone marrow (BM) dysplasia, failure of hematopoiesis and variable risk of progression to AML. One out of three patients diagnosed with MDS will progress to AML, characterized by an increased percentage of myeloid blasts over 20% in the BM. Due to the inherent risk to develop MDS and AML, the BM of patients with FA should regularly be screened for detection of malignant clones.

Most common BM screening techniques include GTG karyotype and fluorescent *in situ* hybridization (FISH). The markers with the highest predictive value for MDS and AML progression in FA include duplication of chromosome 3q (3q+), deletion of chromosome 7q (7q-) or loss of whole chromosome 7 (-7)^4^. Of note, these cytogenetic alterations are associated to MDS with a high risk to progress to AML (high-risk MDS), and therefore high-risk MDS and AML with MDS-related changes are considered to represent a continuum of the same disease.

Detection of MDS and AML clones might be an indication for BM transplantation (BMT) in patients with FA, therefore timely detection of malignant clones is critical for expediting the BMT preparatory regime. BM karyotype and FISH are highly reliable but time-consuming techniques for the detection of malignant clones. For patients at the highest risk of neoplastic transformation, such as patients with FA, the earliest possible detection of abnormal clones is of the utmost importance since rapid evolution of malignant clones in these patients is commonly observed.

Recent single cell RNA sequencing (scRNAseq) technologies have increased our understanding of the transcriptional program of multiple cancer types at unicellular resolution. Specifically, publicly available MDS and AML datasets have been published^5,6^, that can be used to understand the spectrum of the MDS-AML myeloid malignancies.

Machine learning is gaining relevance for analyzing large and complex datasets with large number of features. In the field of machine learning multiple algorithms are available for fitting predictive models to data or for identifying informative groupings within data^7^. Artificial neural networks consist of interconnected nodes that mimic the neuronal connectivity observed in biological brains. Each node, situated within a layer, symbolizes a computed value derived from the preceding layer. These interconnections, referred to as edges, facilitate the transmission of signals across the network’s input, hidden, and output layers^7^

The above mentioned large and complex cancer scRNAseq datasets can be used for training machine learning algorithms with the task to predict malignant cells from complex cell populations, such as the BM of cancer predisposition syndromes, including the IBMFS.

In this work we leverage on multi-layer deep artificial neural networks (DNN) for predicting and identifying single cells with transcriptional profiles similar to MDS and AML in the heterogeneous scRNAseq datasets from the BM of patients with FA. The predicted myeloid malignant cells were found enriched in the LMPP and GMP hematopoietic compartments and have gene expression profiles compatible with malignancy. Here we analyzed the gene expression profile of these predicted malignant cells, propose some of its markers and potentially actionable targets.

## METHODS

### Classifier

#### Data formatting

Given that the training datasets for both classifiers were obtained from different sources^5,6,8^, prior homogenization of the data along with formatting was required. First, in order to train and test the model, two dataset combinations, accounting for the two classifiers, were made using the scRNAseq files. To build the AML classifier healthy donors^8^ and healthy donors and AML patients^6^ were used. Meanwhile, for the MDS classifier, healthy donors^8^ and healthy donors and MDS patients^5^ were used. For each dataset combination a list of intersecting genes was obtained. These shared genes were used as the dataset variables taken as input in the model.

All the combined datasets were divided into two different files, one containing the normalized (CPM), log-transformed (log1p) RNA counts, while in the other the cell annotations corresponding to cell type (e.g. HSPC) and source (e.g. FA patient). An annotated data (AnnData) object (Scanpy) was generated with observations as cells and variables as all the profiled genes. To build the training and testing datasets for the model, the combined datasets were categorically encoded to binarize the annotation labels (healthy = 0, malignant = 1) of each. Both the encoded data and the count matrices for each cell were converted into arrays that were randomly divided into the training sets, accounting for 80% of the data, and the testing sets, with the remaining 20% (**Figure 1A and 1B**).

**Figure 1.**
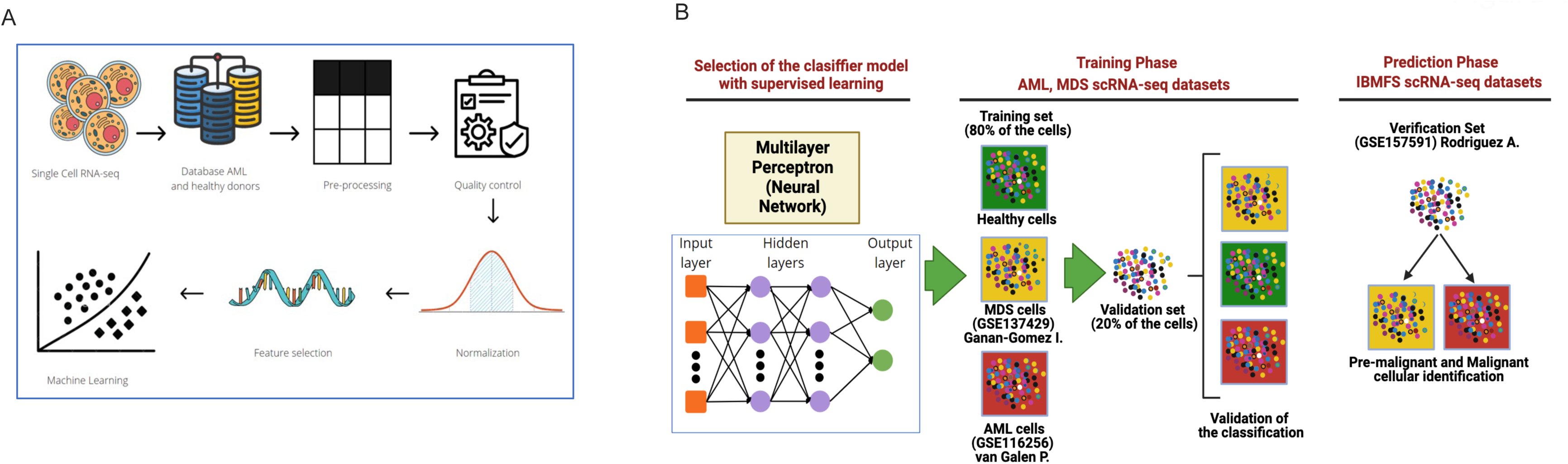
Generation of single cell resolution machine learning-based predictors of malignant myeloid cells. **(A)** Publicly available scRNA-seq datasets of MDS and AML studies were selected for training two machine learning algorithm, one for detection of MDS-related gene expression profiles and a second one for AML related gene expression profiles. Data were pre-processed, homogenized, and normalized before training. **(B)** A multilayer perceptron neural network-based algorithm was selected as the machine learning method. An MDS predictor was trained with 80% of the cells from the Ganán-Gómez et al (2022) dataset (GSE137429); and an AML predictor was trained with 80% of the cells from the van-Galen et al (2019) dataset (GSE116256). 20% of the cells were used for validation of each classifier. Performance of the classifiers were tested in the Rodríguez et al (2021) dataset (GSE157591), where cells with an MDS or AML gene expression profiles were predicted in the HSPCs from patients with FA.

#### Building and testing the model

A deep neural network (DNN) model was built using the deep learning API Keras, that consisted in a sequential layer-based model with a sigmoidal type activation, optimized by Tensorflow stochastic gradient descent method.

#### Predicting malignant cells

Finally, the query datasets, consisting of the FA patients^8^, were formatted as described earlier and their cell types were predicted by using the DNN model. Our model can be found at: https://github.com/BMF-CP-Lab/DNN-AML-MDS-classifier

### Master Regulator Analysis

Master Regulator Analyses (MRA)(10.1038/nature08712, doi:10.1038/msb.2010.31) were performed for the determination of key transcriptional regulators that could be responsible for the establishment of cellular and organismal phenotypes. MRA comparing the transcriptomic regulation of FA predicted-MDS/AML and FA non-MDS/AML cells in the Rodriguez dataset with the CORTO implementation^9^ using a list of all human transcription factors^10^ (TFs). In brief, the analysis aims to identify transcription factors that target differentially expressed genes (DEGs). A transcriptional network is generated based on statistical dependencies between each expression pattern of each TF and the rest of the genes. Each regulon (set of target genes of a TF) is obtained, and the content of DEGs within each regulon is estimated using Gene Set Enrichment Analysis (GSEA) (doi:10.1073/pnas.0506580102). Thus, the TF with the regulon having the highest absolute normalized enrichment score (NES) will be considered as a TF with high potential to be an important regulator of the phenotype. TFs with a positive NES generally upregulate their regulons, while those with a negative NES downregulate them.

### Definition of cell identities in the scRNAseq data sets

The pre-processing and normalization of the Seurat dataset for all samples was conducted as previously described, and the cells were subsequently assigned the cluster number of the "integrated UMAP." Subsequently, the Azimuth algorithm, an automated reference-based approach for single-cell annotation, was applied to the Seurat dataset using the Human - Bone Marrow reference model, which encompasses 297,627 bone marrow cells from 39 donors and 3 different studies^11–13^, and the Human Cell Atlas Immune Cell Census. The cells selected through this annotation-based process were then projected onto the "integrated UMAP-annotated".

### Single cell pathway analysis (SCPA)

The analysis was conducted using the SCPA package, which involved extracting log1p normalized data from each relevant population. The pathways used in the analysis were generated from the publicly available molecular signatures database using the msigdbr package within R. Comparisons were performed using the compare_pathways() function within SCPA, with the only inclusion criteria being gene sets with 15 to 200 genes^14^. Data processing and visualization was carried out using the Seurat, ggplot, and ComplexHeatmap R packages.

## RESULTS

We trained two models based on deep neural networks (DNN) aiming to predict myeloid malignant cells in the BM of patients with FA, specifically cells with transcriptional profiles resembling those of *bona fide* MDS or AML cells. For training we used publicly available scRNAseq datasets (**Figure 1B**) from the BM of patients with MDS or AML, and from healthy donors. For the MDS classifier we obtained a confusion matrix showing a 99% sensitivity, 100% specificity and near to 100% accuracy in MDS-predictions (**Figure 2A**). For the AML classifier, on the other hand, we obtained a confusion matrix showing 98% sensitivity, 94% specificity and 96% accuracy for AML-predictions (**Figure 2B**). Potential overfit of the MDS classifier could be caused by the scarcity of publicly available scRNAseq datasets from MDS, however this issue can easily be corrected in the future with the incorporation of additional datasets from MDS patients.

**Figure 2.**
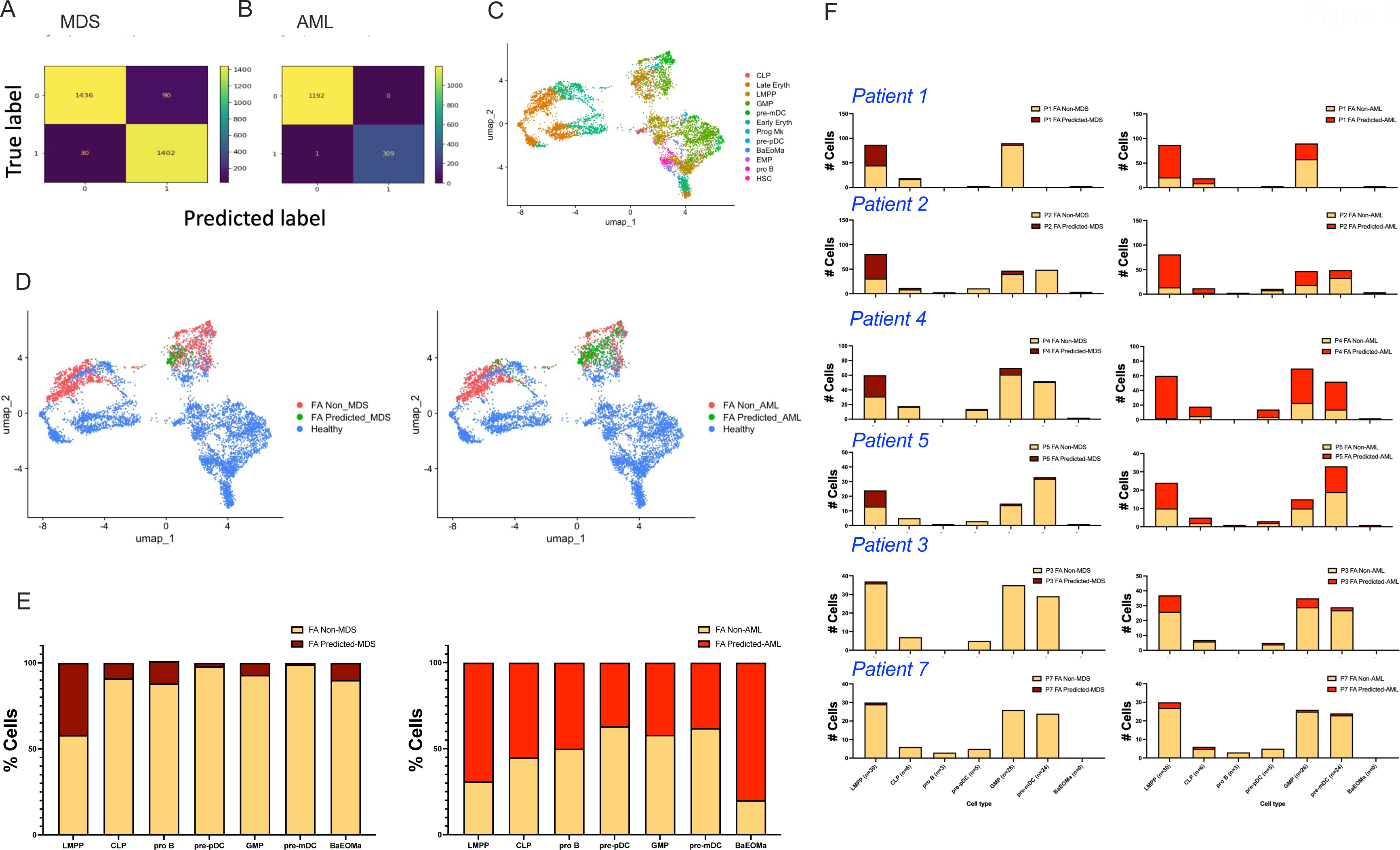
Predicted MDS and AML cells are enriched in the LMPP and GMP compartments of the lymphomyeloid lineage from FA patients. **(A)** Confusion matrix of the MDS predictor showing 99% sensitivity, 100% specificity and 100% accuracy. **(B)** Confusion matrix of the AML predictor showing 98% sensitivity, 94% specificity and 96% accuracy. **(C)** UMAP with cell type annotations of HSPCs from FA patients before MDS and AML predictions **(D)** UMAP with cell type annotations in the Rodríguez et al dataset of HSPCs from FA patients after prediction of MDS cells (*left*) and AML cells (*right*) **(E)** Percentage of cells predicted-MDS cells (*left*) and predicted-AML cells (*right*) in 6 patients with FA. Note the enrichment of MDS and AML cells in the LMPP and GMP compartments. **(F)** Breakdown percentage of the predicted MDS cells (*left*) and predicted AML cells (*right*) per FA patient in the lymphomyeloid compartment. Samples from patients 1, 2, 4 and 5 contain the highest number of cells predicted as MDS and AML. Of note, at examination the bone marrow of patient 1 displayed mildly dysplastic megakaryocytes but no cytogenetically abnormal clones, whereas patient 4 displayed a cytogenetic clone with 7q loss, a well-known marker of clonal haematopoiesis.

Using Azimuth^11–13^ as reference, we annotated the scRNAseq dataset from the BM of patients with FA, and obtained different cell types, including HSC, EMP, LMPP, CLP, GMP, Early Erythroid, Late Erythroid, pro B, pre-pDC, pre-mDC and BaEoMa (**Figure 2C**). Later we visualized the distribution of the cells predicted as MDS (**Figure 2D *left***) and predicted as AML (**Figure 2D *right***) with respect to healthy cells.

Predicted MDS and AML cells were classified on different progenitor types using Azimuth annotations (**Figure 2E**) and identified in every patient (**Figure 2F**). The most common phenotypes of the predicted MDS and AML cells belonged to LMPP, CLP and GMP, corresponding to the cell types where the malignant cell of origin most probably arises in MDS and AML. These results correlate with the clinical characteristics of the patients, an example of this is patient no. 4 whose previous bone marrow cytogenetics detected a clone with deletion of chromosome 7q. Of note, patients no. 3 and no. 7 had the lesser predicted MDS and AML cells among the examined patients.

We explored the Differentially Expressed Genes (DEG) in the predicted MDS and AML cells with respect to the remaining FA cells. In the predicted MDS cells we found increased expression of genes related to inhibitory functions of the immune system, including *CTLA4*, and other genes related to immune functions like *HLA-C*, *IRF5*, which may indicate activation of mechanisms regulating immune functions^15–20^ (**Figure 3A**). In the predicted AML cells, we found increased expression of *WISP3,* which has been found associated to aggressive inflammatory breast cancer and breast cancer with axillary lymph node metastasis, and *CCNA1 (*Cyclin A1) a canonical cyclin that promotes S and G2 phase progression, and that has been reported to be overexpressed in up to 82% of AML cells^15,18,21,22^ (**Figure 3B**)

**Figure 3.**
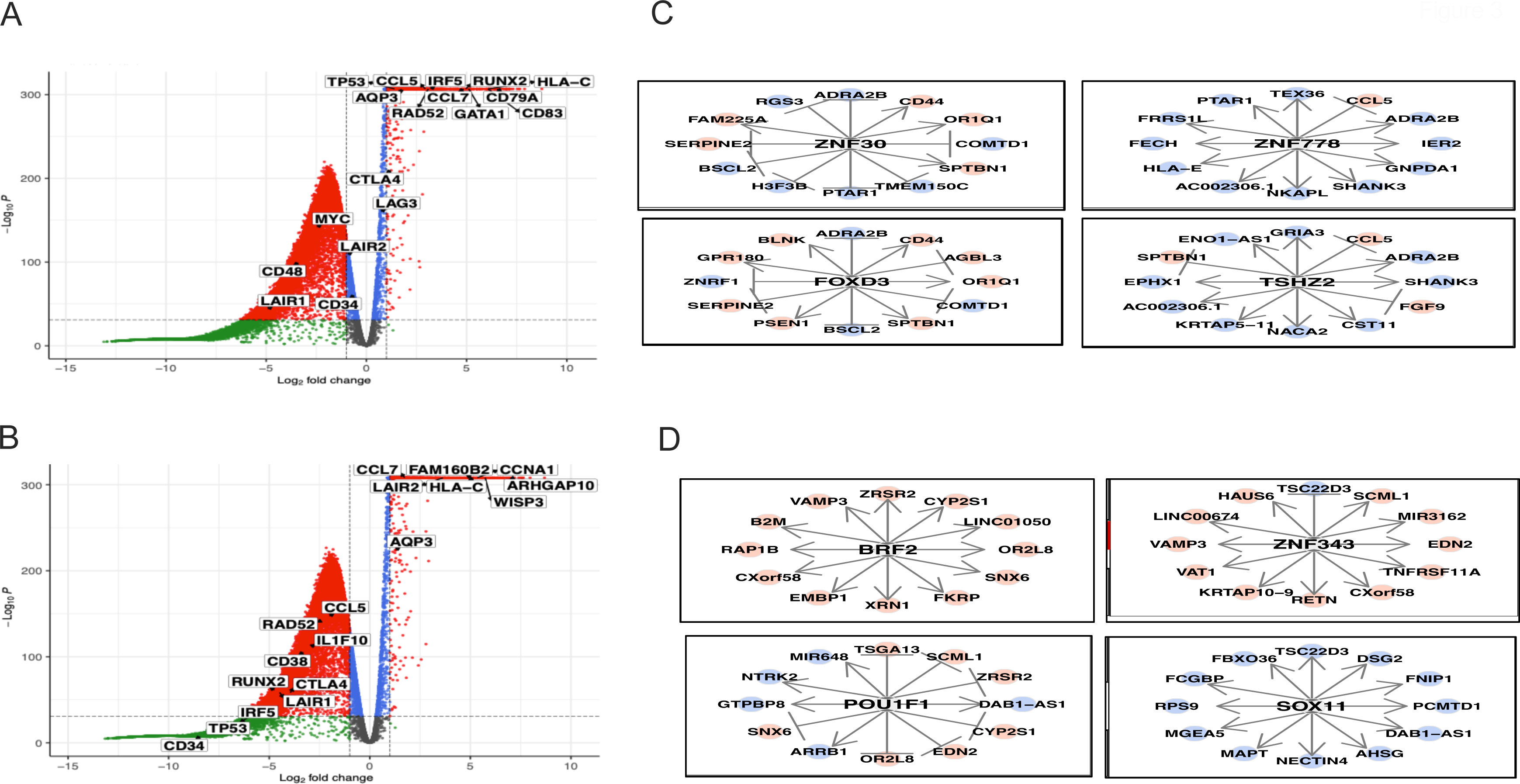
Gene expression profile of the predicted MDS and AML cells from FA patients. **(A)** Volcano plot showing the differentially expressed genes (DEG) in the predicted-MDS cells in comparison to non-MDS cells from FA patients. We observed overexpression of immune inhibitory molecules, such as CTLA4, LAG3 and potential biomarkers like HLA-C, CD83 and AQP3. **(B)** Volcano plot of the DEG in the predicted-AML cells in comparison to non-AML cells from FA patients. Interestingly we observe downregulation of the immune inhibitory molecules (CTLA4 and LAG3), but upregulation of genes related to cancer progression and cellular transformation like WISP3, CNNA1 and FAM160B2. Potential surface markers like AQP3 and HLA-C are shared between the predicted MDS and predicted AML cells from FA patients. **(C)** Master Regulator Analysis (MRA) in the predicted MDS cells from FA patients showing the top 4 transcription factors and their associated genes under transcriptional control. **(D)** MRA in the predicted AML cells from FA patients showing the top 4 transcription factors and their associated genes under transcriptional control.

We performed a Master Regulatory Analysis (MRA)^9^ sought to identify transcription factors and their associated genes that may be overexpressed in the predicted MDS and AML cells. The top 4 transcription factors found to be enriched in MDS included ZNF30, ZNF778, TSHZ2 and FOXD3 which are involved in the regulation of pluripotent Stem Cell Potential, like FOXD3^23^, or regulators of the transcription, like ZNF30, ZNF778 and TSHZ2^24–26^ (**Figure 3C)**. Among the transcription factors enriched in the predicted AML cells we found BRF2, POU1F1 and SOX11, which have been reported in cancer progression or tumorigenic processes^27–29^ (**Figure 3D)**. Although MRA analysis identified important transcriptional factors, MRA does not allow to analyse the specific subsets composing the predicted MDS and AML cells, namely LMPP, CLP and GMP, since it requires larger cell numbers for analysis.

Later we aimed to identify coordinated transcriptional changes in pathways of interest with the single cell pathway Analysis (SCPA) using GO terms (Biological Processes). The pathway “*Negative regulation of cell death*” (gene list shown in the **Supplementary Table 1**) was found to be more active in the predicted MDS and AML cells classified as GMPs (**Figure 4A**).

**Figure 4.**
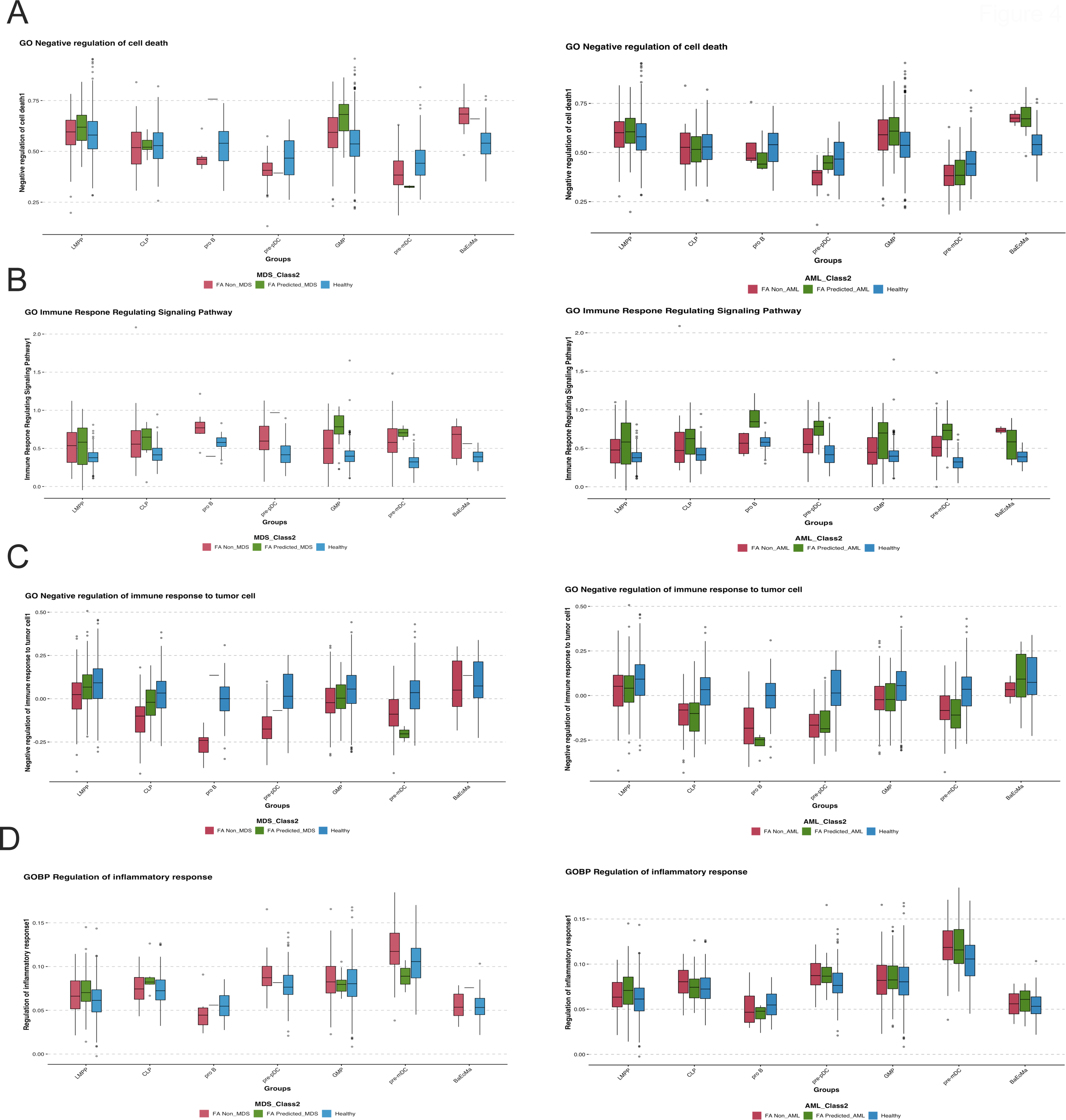
Single cell pathway analysis (SCPA) of the predicted malignant myeloid cells from FA patients shows potential immune modulatory cues. **(A)** SCPA analysis of the GO negative regulation of cell death across annotated cell types in the predicted MDS (*left*) and the predicted AML cells (*right*). **(B)** SCPA analysis of the GO immune response regulating signaling pathway across annotated cell types in the predicted MDS (*left*) and the predicted AML cells (*right*). **(C)** SCPA analysis of the GO negative regulation of immune response to tumor cell across annotated cell types in the predicted MDS (*left*) and the predicted AML cells (*right*). **(D)** SCPA analysis of the GO regulation of inflammatory response across annotated cell types in the predicted MDS (*left*) and the predicted AML cells (*right*).

The pathway “*Immune response regulating signalling pathway”* (gene list shown in the **Supplementary Table 1**), was again found to be more active in the predicted MDS and AML cells classified as GMPs. However, in general the pathway is more active in several cell types present in cells classified as AML, with pro-B, GMP, LMPP and pre-mDC being the main ones. The pathway “*Negative regulation of immune response to tumor cell”* (gene list shown in the **Supplementary Table 1**) was found mostly downregulated in the cells predicted as MDS and AML, which indicates that they have the potential ability to evade the immune system and avoid its elimination.

Finally, modification of the inflammatory environment is one of the most reported processes modified by malignant cells to promote their development and proliferation. We therefore assessed the pathway “*Regulation of the inflammatory response”* (gene list shown in the **Supplementary Table 1**) to evaluate the mechanisms that cells classified as malignant might carry out. In this case, we noticed that the only cells that upregulate this pathway are LMPP, pre-mDC and BaEoMa, classified as AML, while all other cell types have this pathway downregulated (**Figure 4A and 4B lower panels**).

After identifying differential activation of signalling pathways among cell types, we evaluated whether the predicted MDS and AML cells could be communicating or interacting with the other cell types in the scRNAseq dataset of FA patients. Using CellChat^30^ we inferred interactions occurring among cell types, observing that the predicted malignant LMPP and GMP cell types, in both predicted MDS and AML, are the main interactors with other cell types classified as non-malignant (**Figure 5A and 5B**).

**Figure 5.**
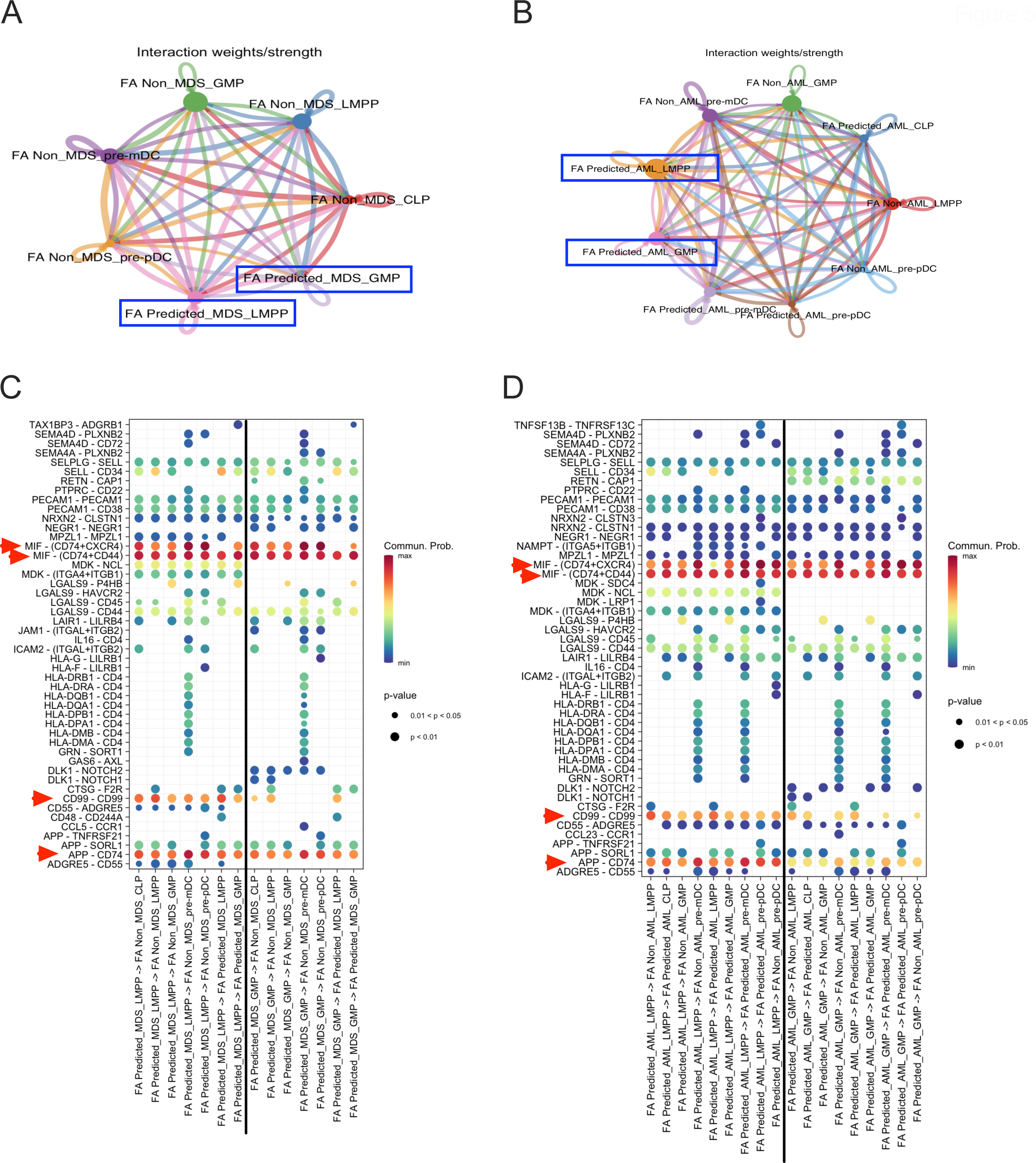
Inference of cellular interactions among the predicted malignant myeloid cells and the non-malignant compartment from FA patients. (**A**) NetVisual circle showing interactions among the predicted MDS cells (LMPP and GMP) with the remaining cell types. (**B**) NetVisual circle showing interactions among the predicted AML cells (LMPP and GMP) with the remaining cell types (**C**) NetVisual Bubble showing the signaling pathways involved in cell interaction among the predicted MDS cells (LMPP and GMP) with the remaining cell types. (**D**) NetVisual Bubble showing the signaling pathways involved in cell interaction among the predicted AML cells (LMPP and GMP) with the remaining cell types.

The most enriched pathways of intercellular communication were the MIF- (CD74+CXCR4) pathway, MIF-CD74 interaction triggers the activation of pro-survival and proliferative Akt and ERK pathways important in tissue reparation^31^, but also this specific interaction has been shown to regulate tumour progression and to determine patient outcomes in advanced melanoma^32^. Another enriched pathway is the CD99-CD99 pathway, a molecule involved in crucial biological processes, including cell adhesion, migration, death, differentiation, diapedesis. CD99 influences processes associated with inflammation, immune responses and cancer, including lymphoma/leukemia^33^ and myeloid malignancies^34^, among others. Finally, the APP-CD74 pathway has been implicated in the production of beta amyloid, but also recent studies have reported this interaction associated to malignant process like melanoma and adenoid cystic carcinoma^35,36^. All together suggests that the predicted malignant cells, mainly LMPP and GMP, early-on activate mechanisms that promote malignancy (**Figure 5C and 5D).**

## DISCUSSION

FA is a rare disease whose prevalence according to the NIH is around 1 out of 136,000 new-borns^37^. This rareness implies that FA does not receive the attention needed from health systems worldwide. In addition, due to a reduced number of patients with FA and their geographical sparsity, obtaining samples from these patients is challenging, leading to limited knowledge about the pathophysiology of this disease, and consequently resulting in their reduced survival.

Assemblage of novel technologies and computational algorithms, however, promises to reduce this breach and help to infer relevant pathophysiological characteristics of rare diseases, such as FA. On the one hand scRNAseq is producing datasets of single cell resolution transcriptional profiles from multiple healthy and diseased tissues, where rare and small populations can be identified^38,39^; on the other hand, multiple machine learning algorithms exist that can be implemented to identify patterns on these datasets, or that can be trained with labelled datasets for identification of transcriptional profiles of interest in inquiry datasets^7,40–42^.

In this work, we sought to make use of machine learning and publicly available scRNAseq datasets to generate a DNN model trained to identify and predict myeloid malignant cells with MDS or AML associated-transcriptional profiles. The later in order to predict the presence of such cells in the BM of six patients with FA for whom BM scRNAseq was previously published^8^.

Our machine learning models based on DNN were independently trained with an MDS scRNAseq dataset^5^ and an AML scRNAseq dataset^6^. Our algorithms were able to predict malignant cells in the BM of the patients with FA, mainly in the LMPP and GMP compartments, which are specifically enriched in the fraction of CD34^+^CD38^-^ cells, thus suggesting a very primitive identity and a transcriptional profile that resembles physiological primitive progenitor cells. Of note, others have proposed that these are important compartments for origin of myeloid malignancy^43^. Interestingly also, our models did not predict myeloid malignant cells in the HSC compartment, which is probably due to the fact that FA patients have very few of these primitive cells, and therefore, the capture of these cells with micro fluidics single cell resolutions technologies was scarce.

Most predicted myeloid malignant cells were detected in 4 out of 6 FA patients. In patient no. 4 previous bone marrow cytogenetics detected a clone with deletion of chromosome 7q. This chromosome abnormality is well-known to have a high negative predictive score^4^, and this sole abnormality places the patients in the high-risk AML group with worst prognosis^44^. Recent work has found that 7q loss is a common event in the carcinogenesis process of FA patients towards AML^45^. In patient no. 1, mildly dysplastic megakaryocytes were detected during routine bone marrow examination, but the bone marrow karyotype was reported as normal. Patients no. 2 and no. 5 were also patients with prediction of MDS and AML cells, however no cytogenetic clones nor morphological changes were detected in their clinical routine. The prediction of MDS and AML cells in these three patients highlights the relevance of searching novel ways to identify malignant progression.

In patients no. 3 and no. 7 a negligible number of malignant cells were predicted, interestingly however, patient no. 3 was previously found to have a clone with chromosome X trisomy, an abnormality that has not been linked to MDS nor AML in FA^4,45^, and might therefore not be of relevance for malignant transformation.

The predicted malignant MDS and AML cells were mainly found in the lymphomyeloid compartment and enriched in the LMPP and GMP subsets. Regrettably not every tool for gene expression analysis supports small numbers of cells, therefore for our initial gene expression analysis we considered the entirety of the predicted MDS or AML cells and compared them against cells from healthy controls and against the FA cells in the same dataset that were not predicted as malignant. This analysis gave us a broad idea about the potential mechanisms dominating the predicted malignant cells.

Our initial results concur with previous reports, where overexpression of *CD74, CTLA-4, HLA-C, CD79A, IRF5* and *LAG3* has been reported in AML and other types of cancer. This indicates their potential participation in tumour development, either as tumour initiators or as immunological checkpoint modulators that will allow malignant progression19,20,31,32,46-49.

Interestingly, the gene expression profile of the predicted MDS cells shows changes related to an active inhibition of the immune system, making us hypothesize that in the MDS stage more immune modulation allows MDS establishment without immune clearance, thus favouring the development of malignant clones^17,19,46^. On the other hand, the expression profile of the predicted AML cells suggests that in these cells the transformation process towards malignancy is already ongoing, since few changes in immune modulation are detected, but more changes in tumour progression molecules seem to take place, including increased expression of *CCNA1*, *HLA-C* and *WISP3*. Importantly these molecules have already been reported as therapeutic targets^16,21,50–52^.

CellChat^30^ is able to predict cell communication with as little as 10 cells, with this tool we were able to specifically detect the interactions of the predicted malignant LMPP and GMP cells with the rest of the cells. The main pathway predicted to mediate communication between the predicted malignant and non-malignant cells is the MIF-CD74 pathway, which is related to cellular repair processes, and has also been reported in some types of cancer such as adenoid cystic carcinoma and melanoma^31,32,35,36^. Communication through the CD99-CD99 pathway was also detected. CD99 has been found to be relevant in lymphoma, leukaemia and myeloid-type malignancies^53,54^. This pathway is particularly interesting since the expression of CD99 in T cells is sought to detect minimal residual disease in acute lymphoblastic leukemia^33^; and some clinical trials are proposing CD99 as a therapeutic target in AML^34,55^.

With our findings we aim to propose a panel of early markers for early identification of myeloid malignant cells in the bone marrow of patients with FA, bearing this in mind we also aim to propose potential therapeutic strategies for targeting these malignant cells before full-blown MDS or AML occurs.

## CONCLUSION

In this work we generated two neural network-based machine learning algorithms using publicly available scRNAseq datasets for the detection of MDS and AML cells. Using these algorithms, we predicted the presence of myeloid malignant cells in scRNAseq datasets from the bone marrow of patients with FA. The predicted myeloid malignant cells were found enriched in the LMPP and GMP hematopoietic compartments and have gene expression profiles compatible with malignancy. Further experimental approaches are sought to confirm the identity of these predicted malignant cells.

## FUNDING

This project was funded to AR through CONAHCYT Ciencia de Frontera 2022, project number 319344 and through PAPIIT-UNAM project IA205022. LAFM receives the scholarship “Estancias Posdoctorales por México 2022 (3)” CVU 407493.

## Notes

### Competing Interest Statement

The authors have declared no competing interest.

