## Supplementary Table 1 for "Prediction of myeloid malignant cells in Fanconi anemia using machine learning"

**SUPPLEMENTARY INFORMATION**

Table 1. Pathways upregulated in MDS or AML malignant cells and their associated genes.

| Pathway Name | Genes involved |
| --- | --- |
| *Negative regulation of cell death* | AIF1, CAV1, CCNG1, CD27, CD34, CD38, CTSH, ENO1, GATA2, HSPA5, HSPB1, HSP90AB1, IL1B, ITGB1, JUN, MYC, NPM1, TKT, RPS3A, RPS6, SOX4, TMBIM6, HSP90B1, XBP1, BTG2, TSC22D1, HERPUD1, PIM2, SLC40A1, VIMP, MYDGF, WNT3A. |
| *Immune response regulating signaling pathway* | *CAV1, CD38, CD79A, CTSH, CYBA, IGHA1, IGHA2, IGHG1, IGHG3, IGHG4, IGKC, IGLC2, IGLC3, IGLL1, OAS1, RPS3, GLTSCR2*. |
| *Negative regulation of immune response to tumor cell* | AHR, IL4I1, CEACAM1, TGFB1, HAVCR2 |
| *Regulation of the inflammatory response* | ADA, CDH5, MIR3909, NR1H3, ADAM8, LPCAT3, TNIP1, CEBPA, CEBPB, CELF1, CTSC, PLK2, DUSP10, USP18, PARK7, MGLL, CHRNA7, NLRP3, CARD16, PGLYRP2, C1QTNF3, IL22RA2, PIK3AP1, LRRK2, CMA1, CCR7, NLRP13, NLRP8, NLRP5, CNR1, CD200R1, ADORA1, ADORA2A, ADORA2B, SIRPA, M APK14, LACC1, LRFN5, C2CD4A, NLRP4, CYLD, CYP19A1, DDT, DHX9, NLRP6, AGER, DNASE1, DNASE1L3, AGT, AGTR1, EDNRB, AHR, AHSG, NLRC3, ELANE, NLRP7, ELF4, TRIM65, NLRP11, ESR1, ETS1, EXTL3, F12, FABP4, FANCA, FANCD2, FCGR1A, FCGR2B, DAGLB, NAPEPLD, FGR, NLRP1, SBNO2, FOXF1, SYT11, UFL1, BRD4, FPR2, MKRN2, SHPK, IL17RA, ALOX5, ALOX5AP, DEFB114, ALOX15, FUT7, RICTOR, FYN, RHBDD3, FBXL2, LETMD1, ABHD12, STAP1, GATA3, PLA2G2D, NUPR1, GGT1, GGT3P, GHSR, FOXP1, IL37, PDCD4, SMPDL3B, RABGEF1, STK39, GPR4, NEAT1, GPR17, METRNL, CXCL17, GPER1, GPR31, FFAR3, FFAR2, GPS2, GPX1, GRN, GIT1, DROSHA, PYCARD, GSTP1, ANXA1, HCK, HGF, HLA-DRB1, AOAH, HLA-E, APCS, BIRC2, BIRC3, XIAP, APOS1, TNC, NLRP9, NLRP10, NLRP14, FFAR4, IFI35, CD200R1L, IGM, IFNG, IGF1, APOE, APE, APP, LILRA5, IL1B, IL1R1, IL2 IL4, IL6, IL6ST, IL10, IL10RA, IL12B, IL13, IL15, IL16, IL17A, IL18, IDO1, INS , IRF3, ISL1, JAK2, KLKB1, KRT1, C2CD4B, LBP, LDLR, LGALS1, LGALS2, LPL, LTA, ARNT, DUOXA2, LYN, MIR138-1, MIR142, MIR15B, MIR181B1, MIR181C, MIR187, MIR223, SMAD3, MAS1, MDK, MEFV, MGST2, MMP3, MMP8, MMP9, CD200, ABCC1, MVK, MYD88, NAIP, ATM, NFKB1, NFKBIA, NINJ1, NKG7, NPY5R, NT5E , OSM, FURIN, PLA2G3, SERPINE1, IL20, IL22, REG3A, FOXP3, TLR7, PDE2A, IL23A, ENPP3, GPRC5B, GHRL, PIK3CG, PLA2G2A, PLCG1, PLCG2, IL20RB, ACP5, SETD4, TLR9, TREM2, PPARA , PPARD, PPARG, CAMK2N1, FEM1A, NLRP2, VPS35, PRKCD, ASH1L, PBK, MAPK7, MAPK13, PROC, MMP26, SUCNR1, PSMA1, SLAMF8, PSMA6, PSMB4, PTGER3, PTGER4, PTGIS, PTGS2, PTPN2 , PTPN6, HAMP, PTPRC, WFDC1, NLRC4, SNX6, IL22RA1, IL21, RB1, ACE2, TRPV4, RELA, BCL6, RORA, BCR, RPS19, S100A8, S100A9, S100A12, SAA1, CCL1, CCL3, CCL5, CCL24, XCL1 , CX3CL1, GPSM3, SELE, SLC39A8, NOD2, NFKBIZ, FNDC4, LRRC19, NCF1, SNCA, SOD1, SPN, SRC, STAT3, STAT5B, BST1, VAMP2, VAMP7, SYK, MIR590, BTK, TEK, TGFB1, TLR2, TLR3 , TLR4, TMSB4X, TNF, TNFAIP3, TNFAIP6, TNFRSF1A, TNFRSF1B, C3, MIR657, TNFSF4, CCR2, TYRO3, ACOD1, SCGB1A1, DAGLA, WNT5A, YES1, ZFP36, MIR766, ZP3, PLA2G7, TNFAIP8L2, MCPH1, RHBDF2 , NLRX1, GSDMD, SETD6, TRIM45, AKNA, NDFIP1, FXR1, ZBP1, ADAMTS12, SHARPIN, BAP1, CASP1, CASP4, PLA2G10, TTBK1, CREB3L3, SEMA7A, CST7, TSLP, TNFSF11, VAMP8, HYAL2, TRADD, SNX4, RIPK1 , TNFRSF11A, IL1RL2, IER3, SPHK1, SIGLEC10, PGLYRP1, TNFSF18, SOCS3, OTULIN, DUOXA1, IL33, NMI, NLRP12, IL1RL1, OSMR, MFHAS1, KLF4, VAMP3, ADIPOQ, CD28, AIM2, PTGES, NR1D1 , CLOCK, CD47, SOCS5, CD81, SPATA2, AREL1, NR1H4, NR1D2 |
